# Immobilized neuroglobin scavenges carbon monoxide from circulating carboxyhemoglobin

**DOI:** 10.64898/2026.04.20.719754

**Authors:** Madison Butler, XueYin N. Huang, Ryan A. Orizondo, Jason J. Rose, Mark T. Gladwin, Nahmah Kim-Campbell, William J. Federspiel, Jesús Tejero

## Abstract

Carbon monoxide (CO) poisoning is responsible for around 50,000 emergency department visits per year in the U.S. alone. With the present standard of care, persistent neurological sequelae occur in ∼30-40% of severe CO poisoning cases. Currently, there is no available targeted molecular antidote for CO poisoning. In previous work, we have developed an antidotal therapy for CO poisoning based on an engineered hemeprotein, human neuroglobin (Ngb-H64Q-CCC). Intravenous infusion of Ngb-H64Q-CCC removes CO from the circulating red blood cells and improves survival in a lethal CO-poisoning mouse model. However, the infusion of heme-containing proteins has inherent heme toxicity risks that may limit the dose that can be used safely without liver or kidney toxicity.

In order to overcome these problems, we have investigated the development of immobilized Ngb in a solid matrix. This approach allows for the development of a CO removal system using an extracorporeal blood circulating system coupled with a stationary matrix with immobilized Ngb-H64Q-CCC. Such system avoids drug infusion and possible organ injury, allows for antidote recycling, and provides advantages for storage and handling of the antidote.

By assessing the efficacy of Ngb-H64Q-CCC immobilized through different linkage strategies, we have identified N-hydroxysuccinimide agarose resin as a viable stationary phase. The immobilized protein shows preserved heme redox activity, can be chemically reduced/oxidized for activation/CO release purposes, and retains its CO removal capacity after successive regeneration cycles. We expect that this novel approach will advance the development of new scavenger-based therapies for CO poisoning.

## INTRODUCTION

Carbon monoxide (CO), generated by the incomplete combustion of organic matter, is one of the oldest poisoning agents known to man [1]. The basic mechanisms of CO toxicity - the replacement of oxygen bound to hemoglobin (Hb) by a CO molecule - have been known for almost two centuries [2, 3]. However, an antidotal therapy for CO poisoning remains elusive. Current therapy is based on the inhalation of 100% oxygen, either normobaric (1 atm), or if possible, at hyperbaric pressures (1.5-3 atm) [4]. The higher oxygen levels enhance the rate of plasma-dissolved CO exchange in the lungs and decrease the carboxyhemoglobin (COHb) half-life from 320 minutes in air to 74 minutes in normobaric 100% oxygen and 20 minutes in hyperbaric (3 atm) oxygen ([4] and references therein). Still, accidental severe CO poisoning cases present a mortality rate around 1-3%, and a high number of survivors (30-40%) develop significant neurocognitive deficits [5, 6]. Overall mortality of CO poisoned patients after the event is about two-fold compared with control subjects [7–9]. Current treatment options remain limited, which highlights an opportunity for improved therapies for this major unmet need.

The binding of CO to Hb is reversible, with dissociation rates of 0.09-0.009 s^−1^ depending on the allosteric state of Hb [10, 11]. This prompted us to consider that CO scavenging would be feasible if a molecule can bind CO with an affinity sufficiently higher than Hb [12, 13]. We have pioneered the study of heme-based scavengers as CO traps that can bind CO in plasma with high affinity [12, 14–16]. In our previous work we have characterized an engineered version of human neuroglobin (Ngb), a recently characterized heme protein [17–20]. This engineered Ngb (Ngb-H64Q-CCC) carries the H64Q/C46G/C55S/C120S mutations and shows extremely high CO binding affinity (over 600-fold higher than human Hb [12]). Intravenous infusion of Ngb-H64Q-CCC can improve survival in poisoned animals in a mouse model of lethal CO poisoning, as well as significantly decrease %COHb in non-lethal models [12]. Ngb-H64Q-CCC also improves mitochondrial function, reverting the inactivation of the complex IV in the mitochondrial electron transfer chain [16]. However, heme proteins in plasma have historically been associated with severe toxicities [21, 22], so Ngb-H64Q-CCC, as a heme protein, would be anticipated to have dosage limitations as well as potential adverse effects [12].

Extracorporeal therapies, such as extracorporeal membrane oxygenation (ECMO), extracorporeal carbon dioxide (ECCO_2_R) systems, and renal replacement therapies (RRT), provide mechanical support when cardiac, respiratory, and/or renal function are compromised [23–25]. These therapies function by utilizing pumps to circulate blood outside the body through membrane systems for gas exchange and solute or fluid removal. Both pulmonary and veno-venous ECMO removal of CO through direct phototherapy have been shown to improve CO elimination in animal studies [26, 27].

Thus, we considered that a high affinity, CO removal device could be applied to a low-flow state standard hemodialysis circuit, like the classic charcoal hemoperfusion methodology [28] or low flow ECCO_2_R systems [29]. To address the concerns about infusing large quantities of circulating heme proteins, we investigated the use of protein immobilization techniques to develop an immobilized Ngb-H64Q-CCC matrix that could be coupled with an *in vitro* extracorporeal circuit to explore the effect of immobilized Ngb-H64Q-CCC resins in conditions simulating extracorporeal CO removal.

## MATERIALS AND METHODS

### Reagents and chemicals

General reagents were obtained from commercial sources without further processing.

### Protein expression and purification

#### Neuroglobin expression and purification

Ngb-H64Q-CCC was purified using standard protocols developed in our lab [12, 16] using *E. coli* SoluBL21 cells (Genlantis) carrying a pET-28 plasmid (Novagen) with the Ngb-H64Q-CCC gene. The protein was purified by sequential DEAE-anion exchange and gel filtration columns (Sephacryl-200HR) as described [12, 16, 30]. Finally, protein samples were concentrated, frozen, and stored at −80°C. UV-Visible spectroscopy and SDS–PAGE were used as purity criteria.

#### Hemoglobin preparation

In order to prepare the human Hb for the efficacy studies, we purified Hb from expired blood units by procedures commonly used in our lab [12, 15, 31–33]. Red blood cells (RBCs) were purified by washing 50 ml of packed RBCs from an expired blood unit with PBS three times by centrifugation at 4000×*g* for 10 min. Then the RBCs were lysed hypotonically by addition of 2-3 volumes of distilled water, and the membrane debris were removed by centrifugation for 30 min at 25000×*g* [32, 34, 35]. The supernatant contains >99% pure Hb [31].

#### Protein concentration and determination of Hb species

The spectral signature of the different Hb species allows for the determination of the individual species by UV-Visible spectroscopy. The concentration of total Hb and the fraction of each oxidation/ligand state (Fe^2+^, Fe^2+^-O_2_, Fe^2+^-CO, Fe^3+^) was determined by UV-Visible spectroscopy and spectral deconvolution using standard spectra as described previously [20, 32, 35, 36].

#### Preparation of COHb

To generate COHb, pure CO gas was passed for 5 min through the head space of a 15 ml tube containing 5-10 ml of Hb (110-150 μM). This was enough to fully saturate the Hb to 100% COHb. As some excess CO remains dissolved in the solution and not Hb-bound, this could alter the observed results by binding to the immobilized Ngb and decreasing its capacity to clear CO from the circulating COHb. To minimize any excess CO in the solution, we mixed the sample with a small amount of oxyHb (1-2ml). The oxyHb bound the remaining dissolved CO in solution and was added until the final COHb concentration was in the 60-90% range. This resulted in final Hb concentrations in the 90-120 μM range. As the affinity of Hb towards CO is very high (K_A_ = 6.0 × 10^8^) [12], a COHb level under 100% ensures that the amount of dissolved CO in the solution is negligible.

### Protein immobilization

#### N-hydroxysuccinimide (NHS)-activated agarose resin

(Cytiva, Marlborough, MA). The resin was washed, swelled in ice-cold 1 mM HCl, and coupled with Ngb-H64Q-CCC in phosphate buffered saline (PBS), pH 7.4, for 24 hours at room temperature. Remaining active sites were blocked with Tris-HCl (pH 8.5) for 2 hours. Nonspecific ligands were washed off with 0.1 M sodium acetate, 0.5 M NaCl buffer, pH 4.5, and the resin was then stored in PBS at 4°C.

#### Carbonyl diimidazole (CDI)-activated agarose

(Cytiva, Marlborough, MA). The agarose storage solution was washed off and replaced with MES buffer (pH 4.7), to which the Ngb-H64Q-CCC was added and left to react for 24 hours at room temperature. Active sites were then blocked by washing with Tris-HCl buffer (pH 8.5) [37] and the resin was then stored in PBS at 4°C.

#### Cyanogen bromide (CNBr)-activated agarose resin

(Cytiva, Marlborough, MA). The resin was rinsed of its storage solution, swelled in cold, acidic conditions (1 mM HCl), rinsed again, then transferred to an alkaline buffer (0.1 M sodium carbonate containing 1 M ethylenediamine). Ngb-H64Q-CCC suspended in 10% DMSO in sodium carbonate buffer was added and left to react for 24 hours at room temperature. The resin was washed and the remaining active sites blocked by washing with Tris-HCl buffer pH 8.5 [38, 39]. The resin was then stored in PBS at 4°C.

#### Circuit setup for COHb scavenging tests

We used a 10ml polypropylene column as main part of the closed circuit to simulate CO scavenging in vivo. The column contains 1ml of immobilized Ngb-H64Q-CCC resin and about 10 ml of COHb-containing buffer. The tubing (MasterFlex MFLX06442-14, 1.6 mm internal diameter) is connected to a peristaltic pump Masterflex L/S Digital Miniflex Pump (Avantor, Radnor, PA) and adds about 0.5 ml to the circuit for a total of 10.5 ml of solution (**Figure 1**). Experiments were conducted using a 1.5 ml/min flow, thus taking 7 minutes for the whole solution to run through the column. Samples (≈ 100 μl) were taken every 7 min (0, 7, 14, 21, 28, 35) to determine the CO removal after the whole solution passed 1, 2, 3, 4, or 5 times through the column. Hemoglobin concentration and COHb levels were determined by UV-Visible spectroscopy and spectral deconvolution as described above [20, 32, 35, 36].

**Figure 1.**
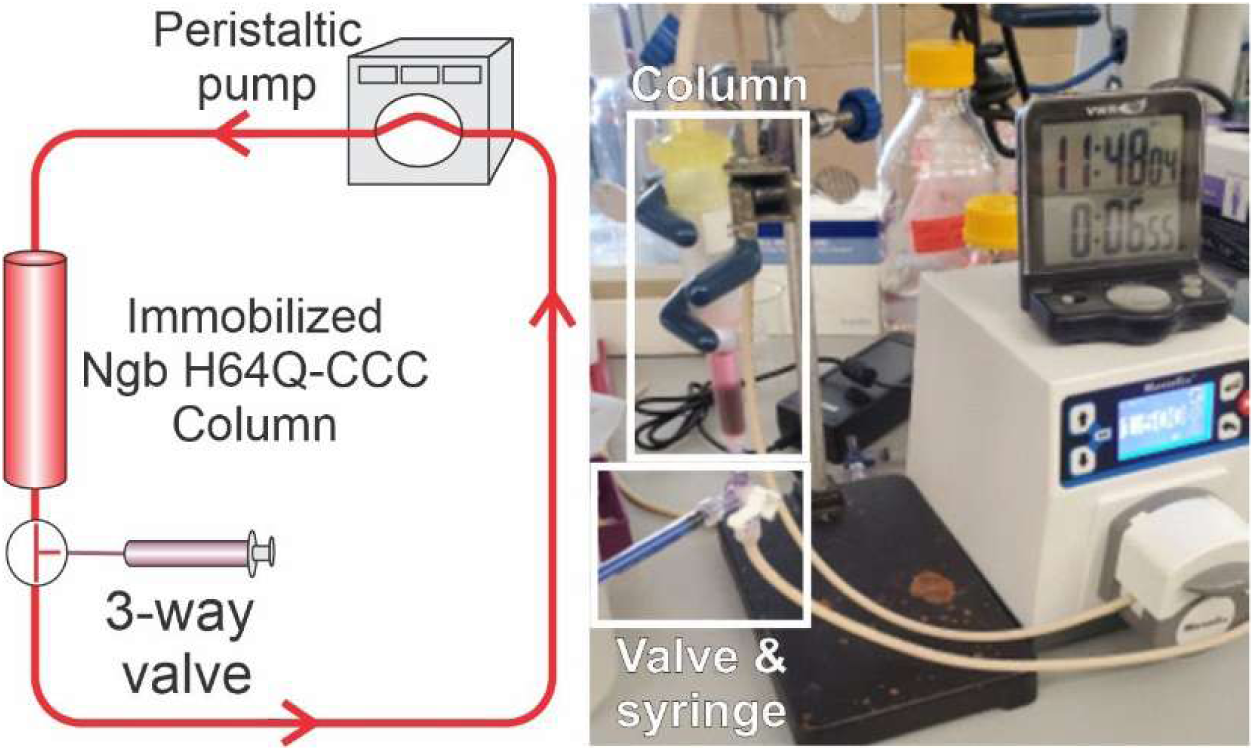
Immobilized Ngb testing setup. Left, scheme of the flow circuit. The final setup comprises a total volume of 10.5 ml where 9.0 are added externally and a 1.5 ml volume is initially PBS (1.0ml over the resin + 0.5 ml of resin void volume). Right, and picture of the setup with the column, 3-way valve and peristaltic pump.

#### Resin activation

The resting state of the Ngb-H64Q-CCC heme has the iron center in its ferric state (Fe^+3^). This oxidation state has minimal affinity towards CO. In order to activate the resin, a small volume of sodium dithionite (≈ 1ml, ≈100 mM) was run through the column. The dithionite was flushed out by running phosphate buffered saline (≈ 10ml). The reduction of the heme iron is easily noticeable by the change in the resin color from dark brown to bright red (**Figure 2**). The ferrous form of the protein is stable enough to be used within ≈1hr (t_1/2_ at 37°C = 50 min [12]).

**Figure 2.**
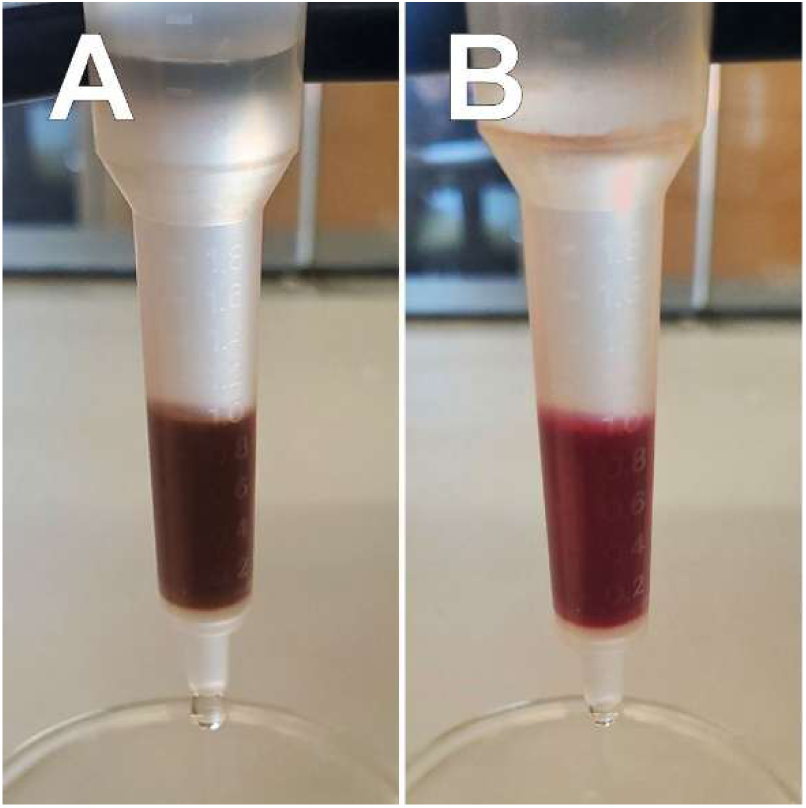
Chemical reduction of the immobilized Neuroglobin shows a noticeable color change. Reduction of the heme center with sodium dithionite leads to a color change of the NHS-immobilized Neuroglobin-H64Q-CCC from the dark brown, ferric heme state (Fe^+3^, Panel A) to the red, ferrous deoxy state (Fe^+2^, Panel B).

#### Resin oxidation/regeneration

After CO scavenging, the protein heme is mainly in the CO-bound state. The affinity of the ferrous Ngb-H64Q-CCC towards CO is very high (*K_A_* = 3.8 × 10^11^ M^−1^) [12]; however as noted above, the ferric heme does not bind CO. Thus, heme oxidizing agents can be used to generate the ferric form, with concomitant release of the bound CO. To fully oxidize the heme group, a small volume of potassium ferricyanide (FeCN) (≈ 1ml, ≈100 mM) was run through the column. Excess FeCN and dissolved CO were flushed out by running phosphate buffered saline (≈ 10ml). After oxidation, the column was stored at 4 °C until further use when it was re-activated with sodium dithionite for CO scavenging.

## RESULTS

We observed different reactivity of the three resins towards Ngb-H64Q-CCC and notable differences in the redox properties of the resin-bound protein. As the heme-bound protein shows noticeable color changes depending on the reduction state of the heme, it is easy to visually detect whether an appreciable amount of the protein remains bound to the resin after the crosslinking reaction and posterior wash steps (**Figure 2**). Firstly, the CDI resins did not bind Ngb in considerable amounts, resulting in an off-white color resin (data not shown) indicating very low Ngb binding and little CO-binding capacity. The CNBr resin did show some binding (**Figure 3A-B**) but there was no noticeable change in color between Fe^3+^ and Fe^2+^ resin, which was inconsistent with known spectral changes of the Ngb-H64Q-CCC bound heme. After further oxidation/reduction cycles, the resin color became progressively lighter, suggesting loss of the heme cofactor. Finally, the NHS resin did show a higher amount of protein bound, as indicated by the darker color, due to the heme group (**Figure 3C**). Upon reduction, the resin color changes from the dark brown of the Fe^3+^ heme to a more reddish color, consistent with the deoxy (Fe^2+^) heme absorption [12, 20] (**Figure 3D**).This change indicates that the heme group retains its redox properties, necessary for proper CO binding. The color change was observed consistently after several oxidation/reduction cycles, suggesting that the cross-linked protein retains its heme-dependent properties. As the other two resins were deemed not suitable for CO scavenging, further experiments were conducted only with NHS-resin.

**Figure 3.**
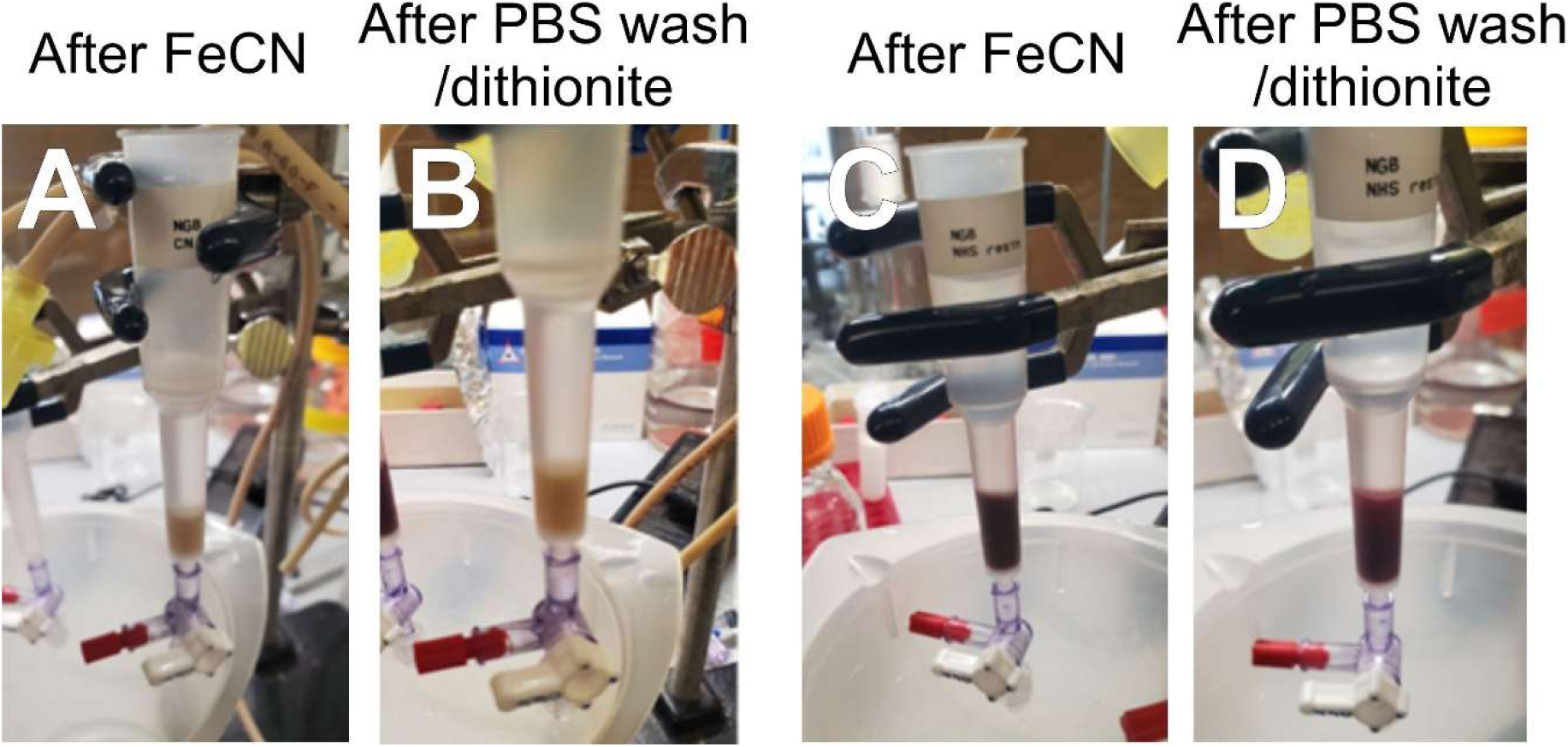
Cross-linked Ngb-H64Q-CCC in CNBr and NHS resins. Panels A and B show the CNBr resin with cross-linked Ngb-H64Q-CCC. Panel A shows the resin after ferricyanide (FeCN) treatment to oxidize all heme iron to Fe^3+^. Panel B shows the same resin after washing the FeCN with phosphate buffered saline (PBS) and addition of sodium dithionite (strong reductant) to reduce the heme iron to Fe^2+^. Panels C and D show the similar process with the NHS-bound protein.

### Carbon monoxide scavenging efficacy

To determine the binding of CO to the immobilized protein, we used a solution of COHb that was circulated through a closed circuit (**Figure 1**). This circuit simulates the arrangement of an extracorporeal blood circulation system, as used in ECMO/ECCO_2_R or dialysis systems [29, 39].

Several tests were conducted to determine the optimal setup (**Figure 1**). After testing different volumes and flow rates we concluded that the most consistent results were obtained with a system containing 10.5 ml of fluid. A flow rate of 1.5 ml/min was determined as optimal.

To determine the amount of CO cleared after passing through the filter, the prepacked column containing immobilized Ngb-H64Q-CCC (≈ 1 ml) was activated by running 1 ml of 10 mM sodium dithionite to reduce the Ngb-H64Q-CCC heme group to the active ferrous (Fe^2+^) state (**Figure 3D**) and excess dithionite was cleared by flushing 5ml of PBS through the column. Then, the column was connected to the circuit and 9.0 ml of COHb of a known initial concentration (90-120 μM) and known %COHb (70-100) was loaded into the column. The circuit was then closed and the pump started at a flow rate of 1.5 ml/min. As the total volume of the circuit (10.5 ml) and flow rate (1.5 ml/min) were known, the whole circuit volume goes through the column every 7 minutes. To represent each of 5 full cycles and monitor the changes in bound CO after every cycle, we took samples (100 μl) at t=0 and then every 7 minutes for 35 minutes. As a control, we used the same resin in the fully oxidized heme state (Fe^3+^), where the heme cannot bind CO, to determine the CO loss due to other factors not related to Ngb scavenging.

A graphic view of the results from the first round of tests is shown in **Figures 4 and 5**. We note that with the inactive resin (Ngb-H64Q-CCC in the ferric state, unable to bind CO), there is a drop in %COHb of around 10% after 5 full cycles (**Figure 4A**). This is consistent with previous observations from our lab and others [12] that COHb decays very slowly in the absence of external scavenging species. However, when the resin was pretreated with sodium dithionite (Ferrous Ngb-H64Q-CCC), the decrease in %COHb was >30% (**Figure 4B**). Replicate experiments showed similar trends. Note that most of the decay in %COHb is observed in the first round, with a less pronounced decay in subsequent cycles (**Figure 4B**). As the decrease in %COHb is dependent on factors like the sample Hb concentrations and initial %COHb, the trends in %COHb do not allow for exact experiment to experiment comparisons. To overcome this limitation, we calculated the amount of CO removed from the solution based on the total Hb concentration, volume and %COHb. The nanomoles of CO scavenged will depend on the available binding sites in Ngb and thus be the same in every experiment independently of the Hb concentrations and %COHb used, as the same Ngb resin was used in all experiments. The trend showing the amount of CO scavenged per cycle for the experiments in **Figure 4** is shown in **Figure 5**. Comparison of the amount of CO scavenged in different experiments (**Table 1**) indicates that the differences between inactive resin (107 ± 27 nmoles) and active resin (269 ± 30 nmoles) are significant (*P*= 0.015, unpaired Student’s t-test).

**Figure 4.**
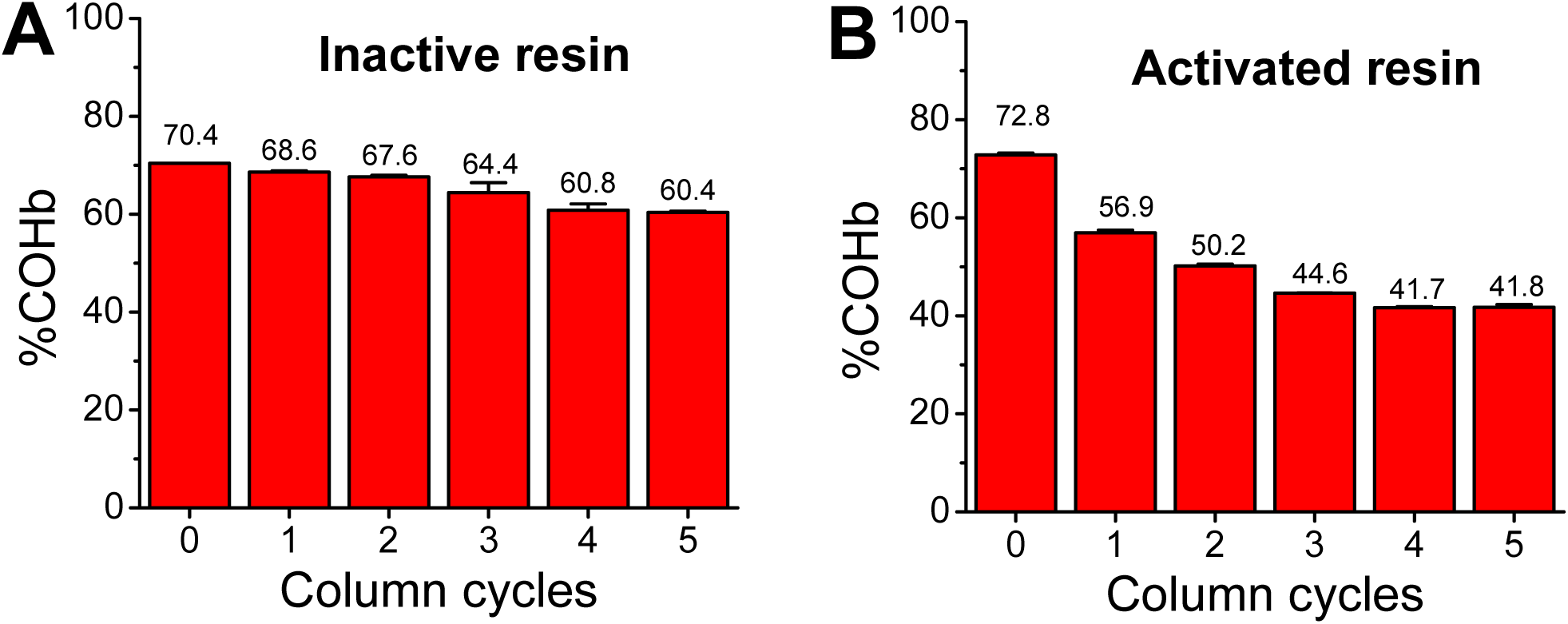
Decrease in %COHb in initial CO scavenging tests. Panel A, results from a single experiment showing the decrease in %COHb of hemoglobin samples after 5 cycles through inactive Ngb-H64Q-CCC-NHS resin. B, observed %COHb from a similar experiment conducted on active Ngb-H64Q-CCC-NHS resin. The number of cycles indicates the circulation of the full circuit volume through the resin (10.5 ml, 7 min at 1.5 ml/min flow). Error bars indicate the standard error of triplicate measurements of the %COHb for each sample.

**Figure 5.**
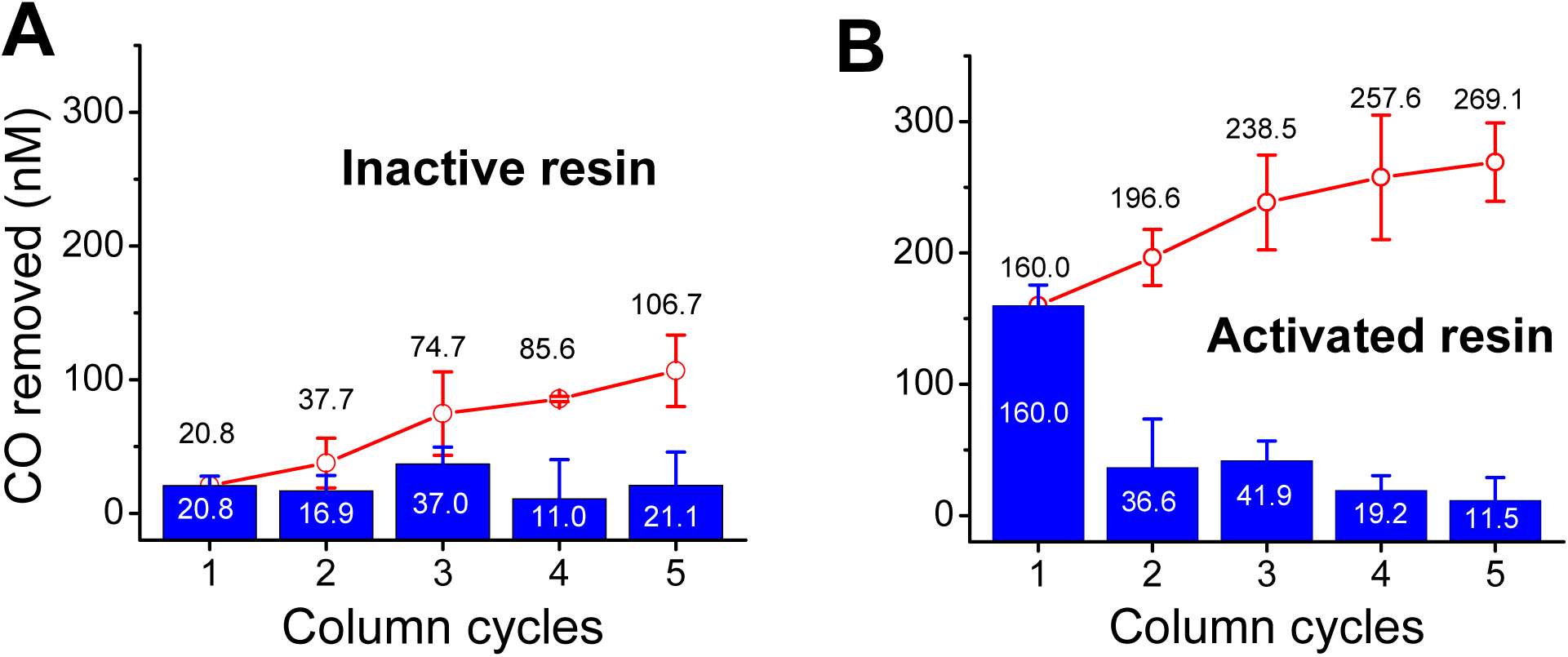
CO removed in initial CO scavenging tests. Panel A, nmoles of CO scavenged per cycle after hemoglobin cycling through inactive Ngb-H64Q-CCC-NHS resin. Panel B, nmoles of CO scavenged per cycle in similar experiments conducted on active Ngb-H64Q-CCC-NHS resin. Blue columns show the amount of CO removed per cycle; the red lines indicate the cumulative amount removed. The number of cycles indicates the circulation of the full circuit volume through the resin (10.5 ml, 7 min at 1.5 ml/min flow).

**Table 1.** Total CO scavenged after 5 column cycles (nmol)

| Initial tests | Inactive resin |  | Active resin |  |
| --- | --- | --- | --- | --- |
|  | 87.7 | Average | 290.2 | Average |
|  | 125.6 | 107 ± 27 | 248.0 | 269 ± 30 |
| ≈75 days later | 28.9 | Average | 239.9 | Average |
|  | 47.3 | 38 ± 13 | 263.1 | 252 ± 16 |
| Average | 72.4 ± 43.2 |  | 260.3 ± 22.1 |  |

In order to study whether the immobilized Ngb-H64Q-CCC retained its binding capacity after long-term storage (stored in the oxidized, ferric form in PBS at 4°C), we conducted similar experiments more than 2 months (75-85 days) after the first experiments.

The results of the experiments to assess resin stability and performance after >5 regeneration cycles are shown in **Figures 6 and 7** and **Table 1**. The same resin was used to evaluate any performance changes after regeneration and storage. The trends observed for the active resin (**Figure 6B**) were similar to those previously seen in the initial round (**Figure 4B**) with a sharp decrease in %COHb in the first cycle and a more modest decrease in later cycles. In the case of the inactive resin, we see a smaller decrease in %COHb (**Figure 6A**) as compared to the decrease observed previously (**Figure 4A**). However, the changes do not reach statistical significance (*P*= 0.063, unpaired Student’s t-test). Notably, the differences in CO scavenging for the activated resin are small (269 ± 30 nmoles in the initial test vs 252 ± 16 nmoles for the second round) and not statistically significant (*P*= 0.28, unpaired Student’s t-test). This observation also suggests that the loss of scavenging activity for the resin is minimal in this ≈ 75 days period. Alternatively, the analysis of the amount of CO scavenged (**Figure 7**) indicates a clear and statistically significant difference between inactive resin (38 ± 13 nmoles) and active resin (252 ± 16 nmoles) (*P*= 0.0029, unpaired

**Figure 6.**
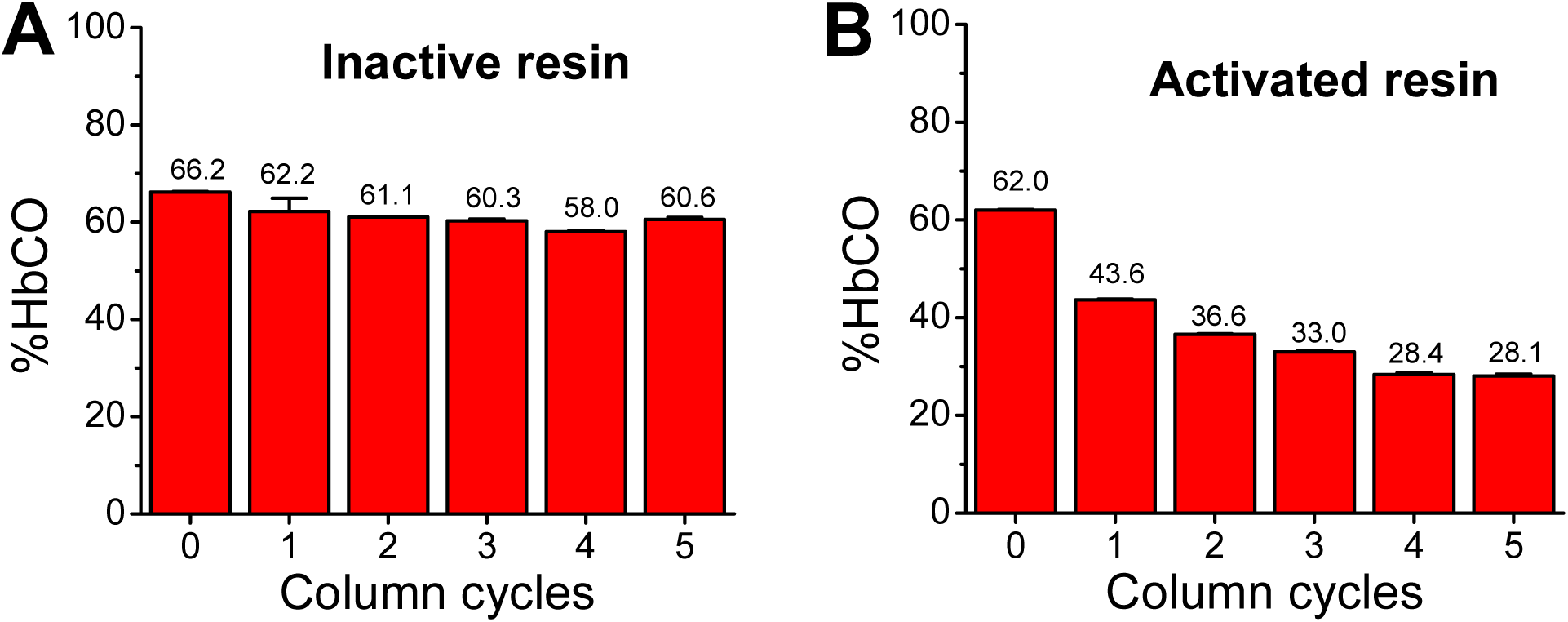
Decrease in %COHb in CO scavenging tests after > 2 month storage. Panel A, results from a single experiment showing the decrease in %COHb of hemoglobin samples after 5 cycles through inactive Ngb-H64Q-CCC-NHS resin. B, observed %COHb from a similar experiment conducted on activated Ngb-H64Q-CCC-NHS resin. The number of cycles indicates the circulation of the full circuit volume through the resin (10.5 ml, 7 min at 1.5 ml/min flow). Error bars indicate the standard error of triplicate measurements of the %COHb for each sample.

**Figure 7.**
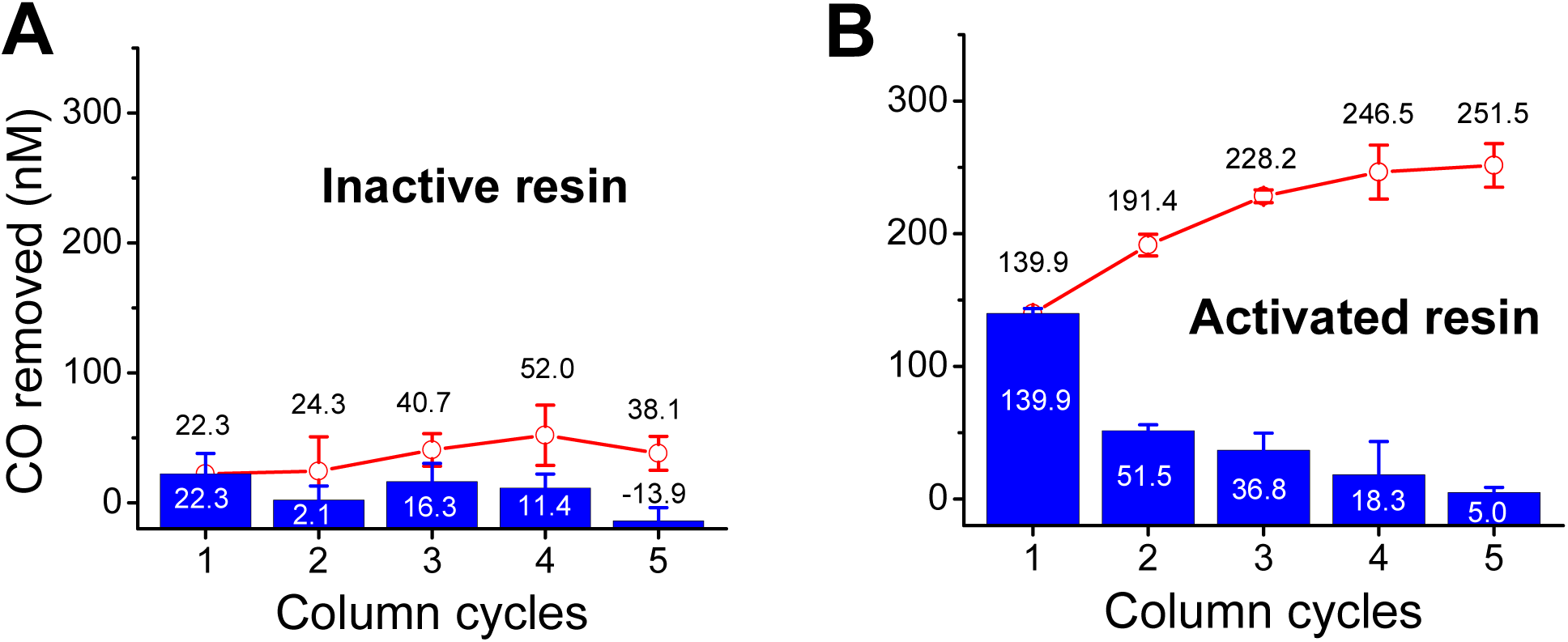
CO removed in COHb scavenging tests after > 2 month resin storage. Panel A, nmoles of CO scavenged per cycle after hemoglobin cycling through inactive Ngb-H64Q-CCC-NHS resin. Panel B, nmoles of CO scavenged per cycle in similar experiments conducted on active Ngb-H64Q-CCC-NHS resin. Blue columns show the amount of CO removed per cycle; the red lines indicate the cumulative amount removed. The number of cycles indicates the circulation of the full circuit volume through the resin (10.5 ml, 7 min at 1.5 ml/min flow).

A summary of the CO scavenging results is shown in **Table 1**. We observed similar values in the two sets of experiments notwithstanding the ≈ 75 day difference between both rounds. The average values indicate a non-specific CO clearance of 72 ± 43 nmol, with a total clearance of 260 ± 22 nmol in the activated resin rounds. and statistically significant difference (*P*= 0.00047, unpaired Student’s t-test)

## DISCUSSION

The development of novel therapies for CO poisoning has advanced significantly in the last 10 years. A few heme protein and porphyrin-based products have been recently studied that show considerable promise and evidence of *in vivo* efficacy [12, 14, 15, 40]. However, there are also challenges in the development of a heme protein-based therapy. Intravenous infusion of high doses of heme proteins presents issues exemplified in the historic efforts to develop a cell-free Hb-based oxygen carrier (HbOC), which have been limited by severe toxicity (renal, hepatic, cardiovascular, thrombotic), thought to be mediated by reactions of the oxygenated heme with nitric oxide (NO) and H_2_O_2_, the formation of superoxide via autoxidation, and the release of heme causing renal toxicity [21, 22]. Ngb-H64Q-CCC is well tolerated in the presence of CO, however some adverse effects were observed when the protein was infused in the absence of CO [12]. A CO scavenger based on the bacterial Regulator of CO metabolism (RcoM) shows improved properties, particularly the lack of hypertensive effect that has hampered HbOC development [14]. Nevertheless, we also recognize that heme toxicity may eventually limit the maximum doses that can be achieved without adverse effects. As the scavenging of CO is stoichiometric, with one molecule of scavenger removing one molecule of CO, large doses of heme-containing antidotes will be ultimately needed for the treatment of patients with high blood COHb levels, requiring robust platforms for protein expression or chemical synthesis to allow the production of significant quantities (grams) of CO antidote. Therefore, approaches that can increase the effective dosage while limiting toxicity are highly desirable.

In this work we have developed an alternative methodology that improves the feasibility of antidotal therapies in two ways. First, by immobilizing the CO antidote onto a resin encased within a fixed extracorporeal column that avoids the need for intravenous infusion and minimizes protein extravasation; and secondly, by allowing for antidote recycling, thereby reducing the antidote production burden. Our data show that the covalent binding of Ngb-H64Q-CCC to an NHS-resin shows optimal properties in terms of protein binding capacity and redox properties of the immobilized protein. We show that the immobilized protein retains its CO-binding capacity after more than 10 regeneration cycles and more than >75 days after immobilization. These results suggest that Ngb-H64Q-CCC cartridges could be stored for an extended period of time and regenerated as needed without significant changes in performance. The ability to visually assess the efficacy of the activation step (**Figure 2**) can also be helpful in practical use. Altogether, we find that this immobilization strategy could be applied to the treatment of CO poisoning with significant advantages over current techniques in development that use intravenous infusion of CO-scavenging molecules. Because it does not enter the systemic circulation, our immobilized heme-based scavenger would minimize possible toxicity issues due to heme-related damage to the liver or kidney. These results are encouraging and provide important proof of concept data for our approach.

Extracorporeal blood circulation therapies, such as ECMO, ECCO_2_R and RRT can provide mechanical life support in conditions of compromised cardiac, pulmonary or kidney function [23–25]. ECCO_2_R and RRT operate through veno-venous configurations at low flow rates that minimize many of the hemodynamic risks associated with ECMO therapy [41, 42]. Our proposed therapy has anticipated compatibility with existing low flow extracorporeal therapies such as extracorporeal carbon dioxide (ECCO2R) and renal replacement therapy (RRT) systems. This would make this therapy more readily available than hyperbaric therapy, which is often logistically very challenging to arrange even in quaternary care centers.

ECMO methods have been investigated in the context of CO poisoning, with some positive results. The use of ECMO combined with hyperoxygenation and phototherapy [26] does increase the rate of CO elimination significantly; other groups have investigated the use of hyperoxygenation alone [43]. Altogether, these methods increase significantly the rate of elimination of CO; although it has been noted that the ECMO can induce oscillations in the hemodynamic pressure [43]. Thus, advances that allow to maintain the CO elimination rate while decreasing the flow rate can improve the feasibility of extracorporeal circulation methods. We show that our antidote can decrease the CO levels on its own, but due to its mode of action we also note that our technology can be combined with oxygenation and/or phototherapy methods, potentially improving efficacy and safety of these approaches.

There are several limitations to this study. Larger volume columns and *in vivo* testing incorporating extracorporeal circulation will be required to understand the efficacy of the immobilized resin in the setting of the dynamic flow and pressure profiles inherent to extracorporeal therapies. The amount of COHb scavenged is expected to increase linearly with the amount of Ngb available, and we expect that scaling up the resin volume and/or the binding capacity will increase the CO scavenging capacity. We do not expect that the resin recycling process and matrix stability will change substantially with scaling up, but this point will need to be reassessed. Future studies in physiologically relevant *in vivo* models will be critical to assess biocompatibility, kinetics, therapeutic effectiveness, and safety profile prior to clinical translation.

## Abbreviations

CDI: Carbonyl diimidazole
CO: carbon monoxide
COHb: carboxyhemoglobin
CNBr: cyanogen bromide
ECCO_2_R: extracorporeal carbon dioxide removal
ECMO: extracorporeal membrane oxygenation
FeCN: potassium ferricyanide
Hb: hemoglobin
Ngb: neuroglobin, Ngb-H64Q-CCC, neuroglobin carrying the H64Q/C46G/C55S/C120S mutations
NHS: N-hydroxysuccinimide
NO: nitric oxide
PBS: phosphate buffered saline
RRT: renal replacement therapies.

## ACKNOWLEDGEMENTS

This work was supported by funding from the National Institutes of Health Grant R01 HL125886 to J.T. and M.T.G., and funding from the Collaboration in HEalth Sciences and EnginERing Startup (CHEERS) Program from the Clinical and Translational Science Institute, University of Pittsburgh, to J.T and W.J.F.

